# No evidence for asymmetric sperm deposition in a species with asymmetric male genitalia

**DOI:** 10.1101/2022.03.19.485007

**Authors:** Sanne van Gammeren, Michael Lang, Martin Rücklin, Menno Schilthuizen

## Abstract

**Background:** Asymmetric genitalia have repeatedly evolved in animals, yet the underlying causes for their evolution are mostly unknown. The fruitfly *Drosophila pachea* has asymmetric external genitalia and an asymmetric phallus with a right-sided gonopore. The complex of female and male genitalia is asymmetrically twisted during copulation and males adopt a right-sided copulation posture on top of the female. We wished to investigate if asymmetric male genital morphology and a twisted gentitalia complex may be associated with differential allocation of sperm into female sperm storage organs.

**Methods:** We examined the internal complex of female and male reproductive organs by micro-computed tomography using Synchrotron X-rays before, during and after copulation. In additon, we monitored sperm aggregation states and timing of sperm transfer during copulation by premature interruption of copulation at different time-points.

**Results:** The asymmetric phallus is located at the most caudal end of the female abdomen during copulation. The female reproductive tract, in particular the oviduct, re-arranges during copulation. It is narrow in virgin females and forms a broad vesicle at 20 min after the start of copulation. Sperm transfer into female sperm storage organs (spermathecae) was only in a minority of examined copulation trials (13 / 64). Also, we found that sperm was mainly transferred early, at 2 - 4 min after the start of copulation. We did not detect a particular pattern of sperm allocation in the left or right spermathecae. Sperm adopted a granular or filamentous aggregation state in the female uterus and spermathecae, respectively.

**Discussion:** No evidence for asymmetric sperm deposition was identified that could be associated with asymmetric genital morphology or twisted complexing of genitalia. Male genital asymmetry may potentially have evolved as a consequence of a complex internal alignment of reproductive organs during copulation in order to optimize low sperm transfer rates.

## Introduction

Animal genitalia are often remarkably complex and reveal a high degree of variation between species (Eberhard, 1985). Sexual selection has been proposed as the chief driver for both the rapid diversification and the great morphological complexity of genital structures through sexual conflict between females and males at different levels of reproduction, including access to copulation, sperm competition inside the female, and control over post-mating sperm storage and usage for fertilization (Thornhill, 1983; Eberhard, 1985; Arnqvist, 1998; Birkhead & Pizzari, 2002; Chapman et al., 2003). In general, genitalia enable ejaculate transfer during copulation from the male into the female and mediate inter-sexual communication during copulation (Eberhard, 1985, 1994).

Asymmetric genitalia are observed in many species and must have recurrently evolved from symmetric ancestors (Huber, Sinclair & Schmitt, 2007; Huber, 2010; Schilthuizen, 2013). They have been associated with mating behaviors (Otronen, 1998; Kamimura, 2006; Huber, 2010) and in particular with specific interlocking of female and male genitalia (Kamimura, 2006; Holwell et al., 2015; Rhebergen et al., 2016; Richmond, Park & Henry, 2016). However, the ultimate evolutionary cause for these transitions from symmetry to asymmetry are not fully understood (Schilthuizen, 2013). To help solve this puzzle, detailed analyses of the function and the internal configuration of asymmetric genitalia during copulation are helpful, as already evaluated in a few cases. In anseriform birds (waterfowl), for example, asymmetric female and male genitalia potentially co-evolved to control fertilization success. The female vagina is coiled clockwise and has dead end sacs that potentially function to avoid insemination via the male phallus, which is coiled in counter-clockwise direction (Brennan et al., 2007). Also, complementary antisymmetric genitalia have been suggested to increase reproductive success in snails of the subgenus *Amphidromus* s. str. The genitalia of these hermaphrodite species are coiled and favor copulation of interchiral pairs that better match for sperm transfer and that circumvent sperm digestion (Schilthuizen et al., 2007; Schilthuizen & Looijestijn, 2009). In insects, the evolution of asymmetric genitalia was proposed to have evolved in response to changes in mating position, potentially affecting the coupling efficiency of female and female genitalia during copulation (Huber, 2010). Asymmetric genital morphology was suggested to optimize these contacts that can be misaligned through a changed mating position. However, no experimental data exist to test this hypothesis and generally little is known about the internal alignments of the intermittent phallus inside the female reproductive tract. Such studies have, however, become feasible in recent years due to the advances in micro-computed tomography (Mattei et al., 2015; Woller & Song, 2017; Dougherty & Simmons, 2017; Gutiérrez et al., 2018).

Males of *Drosophila pachea* have strikingly asymmetric genitalia. They possess external genital lobes with the left lobe being approximately 1.5 times longer than the right lobe (Pitnick & Heed, 1994; Lang & Orgogozo, 2012). In addition, the phallus is asymmetrically bent, harbors ventrally asymmetric spurs and the male gonopore is positioned at the right dorsal tip of the phallus (Acurio et al., 2019). Apart from this asymmetric genital morphology, *D. pachea* also mates in a right-sided copulation posture, with the male antero-posterior midline being shifted about 6-8° to the right side of the female midline (Lang & Orgogozo, 2012; Rhebergen et al., 2016; Acurio et al., 2019). This one-sided mating posture is associated with asymmetric genitalia coupling and an asymmetric twist of the female ovispositor (Rhebergen et al., 2016) potentially resulting in a specific positioning of the male gonopore inside the female reproductive tract.

Sperm storage and usage for fertilization has been associated with sexual conflict and the evolution of sexual characters with respect to sperm selection from multiple copulations, sperm viability and temporal unlocking of copulation and fertilization (Orr & Zuk, 2012; Edward, Stockley & Hosken, 2015). *Drosophila* species store sperm after copulation in two types of sperm storage organs, the tubular seminal receptacle and the paired spherical spermathecae (Pitnick, Markow & Spicer, 1999). Both types of organs are connected to the anterior end of the female uterus. In particular, *D. pachea* and two sister species *D. acanthoptera* and *D. wassermani* were reported to store sperm exclusively in the spermathecae (Pitnick, Markow & Spicer, 1999). Sperm length also varies enormously among *Drosophila* species and range from 0.3 mm in *D. persimilis* to 58 mm in *D. bifurca* (Pitnick, Markow & Spicer, 1995). In particular, *D. pachea* and its sister species *D. nannoptera* also produce giant sperm with lengths of 16.5 and 15.7 mm, respectively (Pitnick & Markow, 1994). Migration capacities of such long sperm require further investigation and it is unknown how giant sperm is organized post copulation inside female reproductive tract.

It is known that in certain Diptera that females preferentially store sperm in specific spermathecae for high-quality males (Otronen, 1998; Hellriegel & Bernasconi, 2000). It is therefore conceivable that the asymmetric male genitalia in *D. pachea* may have evolved to enhance the chances that sperm would be deposited in or near such preferential sperm storage organs. The aim of our study was to investigate if male genital asymmetry results in an asymmetric internal configuration of the copulatory complex in *D. pachea* which may allow sperm to be directed specifically towards one of the female spermathecae. We carried out microcomputed tomography (microCT) of snap-frozen *D. pachea* mating couples before and during copulation to investigate the positioning of reproductive organs as well as ejaculates during copulation. Furthermore, we investigated when during copulation sperm transfer occurs and what aggregation states ejaculate mass assumes during and after transport into the spermathecae.

## Materials & Methods

### *Drosophila pachea* maintenance and virgin collection

*D. pachea* stock 15090-1698.02 from the San Diego *Drosophila* Species Stock Center (now The National Drosophila Species Stock Center, College of Agriculture and Life Science, Cornell University, USA) was maintained in 25 × 95 mm plastic vials containing 10 mL food medium (60 g/L brewer’s yeast, 66.6 g/L cornmeal, 8.6 g/L agar, 5 g/L methyl-4-hydroxybenzoate and 2.5% v/v ethanol). In addition, 40 μL of 5 mg/mL 7-dehydrocholesterol (dissolved in ethanol) was mixed into the food medium with a spatula (standard D. pachea food). Flies were transferred to fresh vials every 2 to 4 days. In order to isolate virgin individuals, adult flies at 0-3 day after emerging from the pupa were anaesthetised on a CO2-pad (INJECT+MATIC Sleeper) under a stereo-microscope Stemi 2000 (Zeiss), separated according to sex and maintained in groups of 20-30 female or male individuals until they reached sexual maturity, about 14 days for males and 4 days for females (Pitnick, 1993).

### micro-CT scanning

Sexually mature virgin couples of *D. pachea* were introduced each into a 2-mL reaction tube (Eppendorf) that was roughened on the inside. Flies were observed at room temperature for at most 1 hour or until copulation had ended. As soon as a couple started to copulate, time was recorded and the couple was snap-frozen by submerging the Eppendorf tube for 20 sec in liquid nitrogen after one of the following intervals: 2 min, 8 min, 15 min, 20 min, and 120 min after the start of copulation. Tubes were filled with cold (−20°C) absolute ethanol and were stored at −20°C for about a week. Finally, the absolute ethanol was replaced by 80% ethanol and samples (mating complexes) were stored at room temperature. A total of 40 samples were prepared for microcomputed tomography scanning (micro-CT) by critical-point drying, including 25 mating complexes, 9 single females 2 hours after copulation start (ACS) and 6 virgin females (Table 1). First, the complexes were fixed by successive 20-min incubations in 90% ethanol, 96% ethanol, and twice in absolute ethanol. Samples were then dried in the Automated Critical Point Dryer EM CPD300 (Leica) (program 1, parameters: ‘CO2 IN’ speed slow, delay 120 sec; ‘Exchange’ speed 5, cycles 12; ‘gas OUT’ heat slow, speed slow). Samples were subsequently mounted on an aluminium stub with the male’s dorsal side (mating complexes) or the female dorsum (for single female specimen) glued to the stub surface. The ventral side of the female pointed upward and was therefore visible. Twelve scans of different mating complexes were performed with the Zeiss Xradia 520 Versa micro-CT scanner at Naturalis Biodiversity Center in order to adjust experimental conditions. Scan 14 was used to build the model of the female reproductive tract (Table S1). A total of 28 single fly specimen or mating complexes (Table S1) were scanned using Synchrotron Radiation X-rays Tomographic Microscopy (SRXTM) at the TOMCAT (X02DA) beamline of the Swiss Light Source, Paul Scherrer Institut, Switzerland. Of those, 26 scans could be used for data analysis (Table 1, Table S1). Absorption and phase contrast scanning modes were performed at 10, 20, and 40-fold magnification and 10 KeV, with 200 ms to 600 ms exposure time (Table S1). Projections were post-processed and rearranged into flat- and darkfield-corrected sinograms, and reconstruction was performed on a Linux PC farm, resulting in isotropic voxel dimensions of 0.65 μm, 033 μm and 0.16 μm.

**Table 1:** Number of scanned *D. pachea* copulation complexes and single females

| Virgin female | Virgin male | Copulation complexes, after copulation start (ACS) |  |  |  | Mated female | Mated male |
| --- | --- | --- | --- | --- | --- | --- | --- |
|  |  | 2 min | 8 min | 15 min | 20 min | 2 h ACS | 2 h ACS |
| 2 | 3 | 5 | 4 | 4 | 3 | 4 | 1 |

### Ejaculate quantification in female spermathecae

In the laboratory, *D. pachea* copulation lasts for about 30 min (Lang & Orgogozo, 2012; Rhebergen et al., 2016; Acurio et al., 2019; Lefèvre et al., 2021). To further examine when during copulation sperm transfer would take place, we interrupted copulation of a single couple by vigorous shaking at various time points after copulation start and examined the presence of ejaculate about 24 h later in the female sperm storage organs (Table S2). We prepared mating complexes as described above and monitored the mating progress by annotation of the courtship and copulation duration. The former was defined as the period from the first male “licking” courtship behavior (Spieth, 1952) to the start of copulation. Copulation start was defined as the moment when the male mounted the female and copulation end was considered to be the moment when the male had dismounted, with genitalia and forelegs being fully detached from the female. Copulation was interrupted at 4 min, 8 min, 12 min, 18 min, and 24 min after copulation start by shaking the tube until the couple separated. The male was stored in 96% ethanol and the female was transferred into a food vial for at least 12 hours at 25°C. The number of eggs laid were counted (Table S2) and the female paired spermathecae were prepared by opening the dorsal side of the abdomen and then separating them from the reproductive tract. The remainder of the female’s body was stored together with the corresponding male in 96% ethanol.

For imaging, spermathecae were transferred into a transparent dissection dish, filled with phosphate buffered saline (PBS, 137 mM NaCl, 2.7 mM KCl, 10 mM Na2HPO4, 1.8 mM KH2PO4, pH 7.4) and examined in lateral view with a VHX2000 (Keyence) microscope equipped with a 100-1000x VH-Z100W (Keyence) zoom objective at 500 fold magnification. For data analysis, file-names were replaced by a three-digit random number so that sperm presence was quantified blindly with respect to the annotated copulation duration. Ejaculate presence or absence in the spermathecae was annotated based on visual inspection according to previously published protocols (Jefferson, 1977; Lefèvre et al., 2021).

Sperm cells inside the female spermathecae of synchrotron X-ray tomographic microscopy samples were quantified by counting sperm cells in three transverse sections that corresponded to 25%, 50%, and 75% of each spermatheca along the axis from the basis to the apical tip (Table S3).

## Results

### The male phallus is asymmetric

The phallus of *D. pachea* has a complex and asymmetric shape (Acurio et al., 2019) (Figure 1). In dorsal/rostral view, the distal part (apex) is rather flat but pointed at the apical tip (Figure 1B). It is strongly curved or folded at the base. The gonopore is positioned dorso-apically on the right side of the aedeagus. In lateral view (Figure 1A, C), a broadened ridge on the apical side and a slender ridge on the more distal side are observed. The ejaculatory duct is located at the right side of the phallus and ends in the right-sided apical gonopore (Figure 1A, B). In contrast, the uterus of a virgin female appearsto be bilaterally symmetric (Figure 2A). It appears to be S-shaped and forms a major sack with a dorsal and a ventral blind protrusion (Figure 2B). The anterior part of the uterus forms a narrow tube that leads to the ovaries (ovaries are not included in the 3D-model).

**Figure 1:**
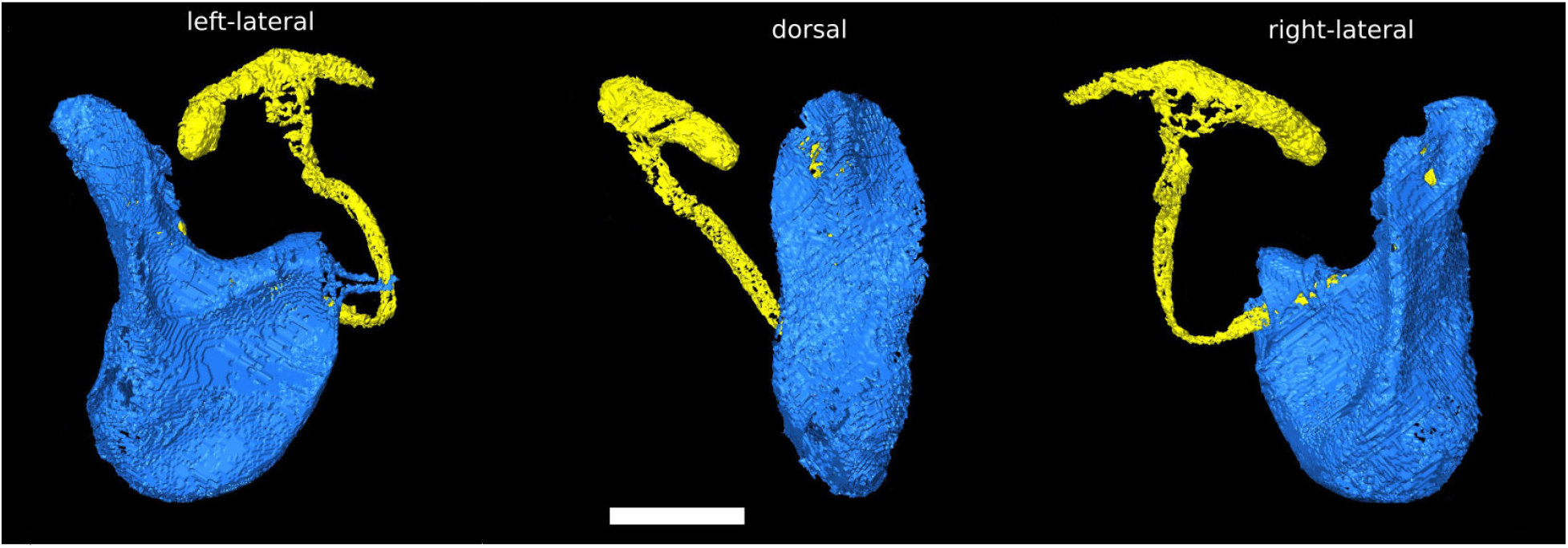
Asymmetric *Drosophila pachea* phallus morphology. 3D-Model of the male phallus (blue) and ejaculatory duct (yellow). The scale bar is 100 μm.

**Figure 2:**
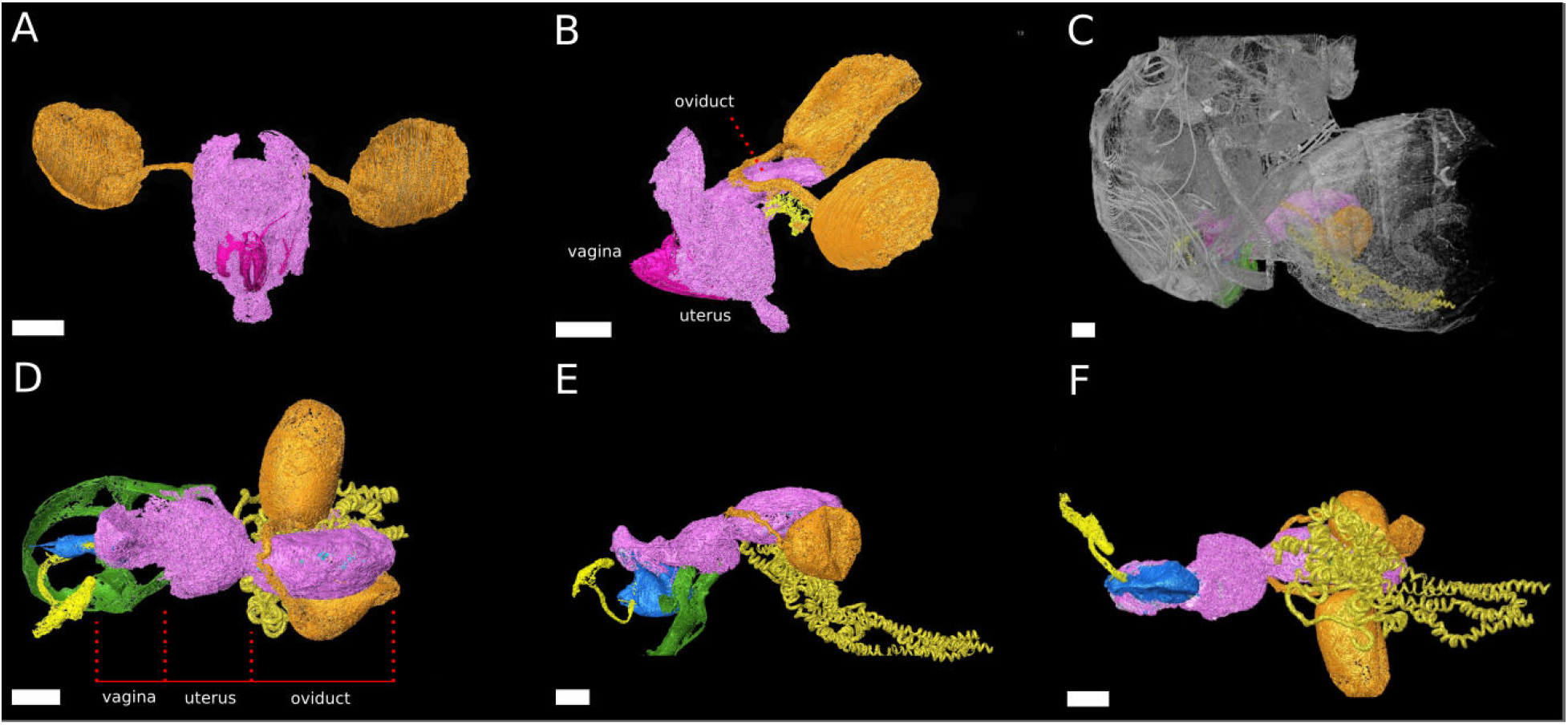
The female reproductive tract re-arranges during copulation. 3D-Models of virgin female *D. pachea* **(A,B)** and copulating couples **(C-F).** The female accessory gands and ovaries are not included into the model, A) Virgin reproductive tract (pink) in caudal view, ovipositor (magenta) and spermathecae with spermathecae ducts (orange). B) Virgin uterus in lateral view (color codes as in A) and the receptaculum (gold). **C)** Copulating couple in lateral view, exterior cuticles of female and male (grey) semi-transparent and relative location of colored the female repoductive tract and male external genitalia (green). **D-F)** Copulation complex of **C)** at higher magnification: female reproductive tract (pink), spermathecae and ducts (orange), receptaculum (gold) and male-phallus (blue), ejaculatory tract (yellow) and genital arch (green), red lines indicate assigned compartment borders of the female reproductive tract into vagina, uterus and oviduct. **D)** Dorsal view, **E)** lateral view, **F)** ventral view. The scale is 100 μm.

### The reproductive tract rearranges during copulation

The uterus re-arranges and re-orients during copulation, similar to previous observations in *D. melanogaster* females (Adams & Wolfner, 2007; Mattei et al., 2015) (Figure 2). The model in Figure 2 describes the configuration of the uterus at 20 min after copulation start. At this time point, the repoductive tract can be divided into three compartments: 1) the vagina, which is the most caudal part, 2) a balloon-shaped medial uterus, and 3) a broadened oviduct (Figure 2D). The vagina appears to be elastic. It harbors and adopts the shape of the dorsal male phallus, which does not reach into the uterus but stretches the vagina in opposite dorsal-caudal direction. The male genital lobes (Figure 2E) are positioned externally against the ventral female abdomen wall (not shown in the model). Together with the phallus they appear to be used to lever or clamp the male abdomen against the female abdomen (Figure 2D, E). The caudal positon of the phallus tip with the male gonopore is most distantly to the anterior parts of the female reproductive tract. Ejaculate needs to pass through the vagina and along the phallus towards the uterus in order to reach the female spermathecae (Figure 2E). The vagina is dorsally pointed at its anterior end where it resembles the curved phallus base. (Figure 3E). The expanded medial uterus is separated from the anterior oviduct by a constriction, which also links the ducts to the spermatheacae and the seminal receptacle at the dorsal and ventral side, respectively (Figure 2D – F). The oviduct forms a broad vesicle of the size of the uterus, whereas it is narrow in virgin females and in *D. melanogaster* (Adams & Wolfner, 2007) (Figure 2B, E) (ovaries were not included in the model).

**Figure 3:**
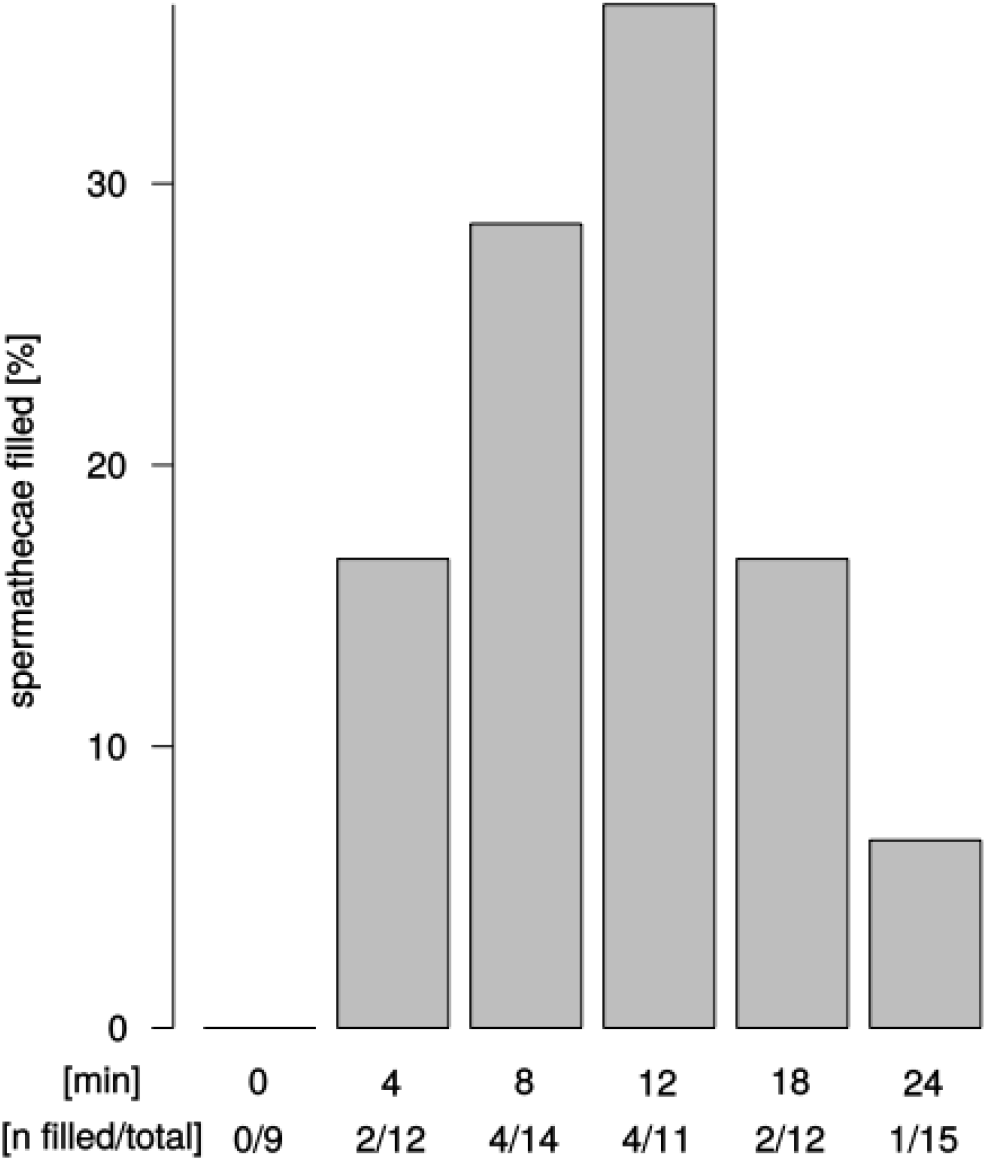
Ejaculate transfer occurs early during copulation. Fraction of female spermathecae that revealed apparent sperm mass from copulations that were interrupted at different time points after copulation start.

### Sperm transfer can occur at 2 min after copulation start and is found in both spermathecae

We observed sperm presence in female spermathecae by dissections of females about 12-24 h after copulation. The spermathecae of virgin females (n=9) were empty. Apparent sperm masses were identified after 4 min of copulation (2/12 trials) (Figure 3). Overall, sperm transfer into the spermathecae occurred only in a minority of trials (13/64). Highest frequencies of ejaculate presence were observed after 8 min and 12 min (4/14 and 4/11 observations, respectively) of copulation. Unexpectedly, longer copulation duration of 18 min and 24 min yielded fewer cases of sperm presence in the spermathecae (2/12 observations and 1/15 observations, respectively). In all 13 trials with observable sperm transfer, both spermathecae revealed apparent sperm mass. In order to quantify different sperm levels in our synchrotron X-ray tomography scans, we counted sperm in a series of 3D-scans (Table S1). Upon copulation, sperm was present in the female reproductive tract in the majority of scans (14/20 scans), but in different parts of the reproductive tract: in the uterus, spermathecae, spermathecal tubes, and in the broadened anterior part of the uterus (Tables S1, S3). Sperm presence in the uterus was even detected in one couple at 2 min after copulation start (1/5 scans) and spermathecae contained sperm at least 8 min after copulation start (1/4 scans) (Table S1). Sperm mass abundance varied between the left and the right spermatheca but no particular bias of sperm storage towards the left or the right spermatheca was observed among couples or at different time points after copulation start (Table S1).

### Different sperm morphologies in the female reproductive tract

Similar to previous findings (Pitnick, Markow & Spicer, 1999), no sperm cells were found in the seminal receptacle. By means of high-quality scans using SRXTM it was possible to visualize the giant sperm cells within the female reproductive tract (Figure 4, Table S2). The sperm cells appear to adopt different aggregation states in different parts of the uterus. They are spherically coiled or granular in the medial uterus (Figure 4A), while they are filamentous and potentially consisting of multiple sperm cells when entering the spermathecae (Figure 5B). No distinction could be made between the sperm tails and heads.

**Figure 4:**
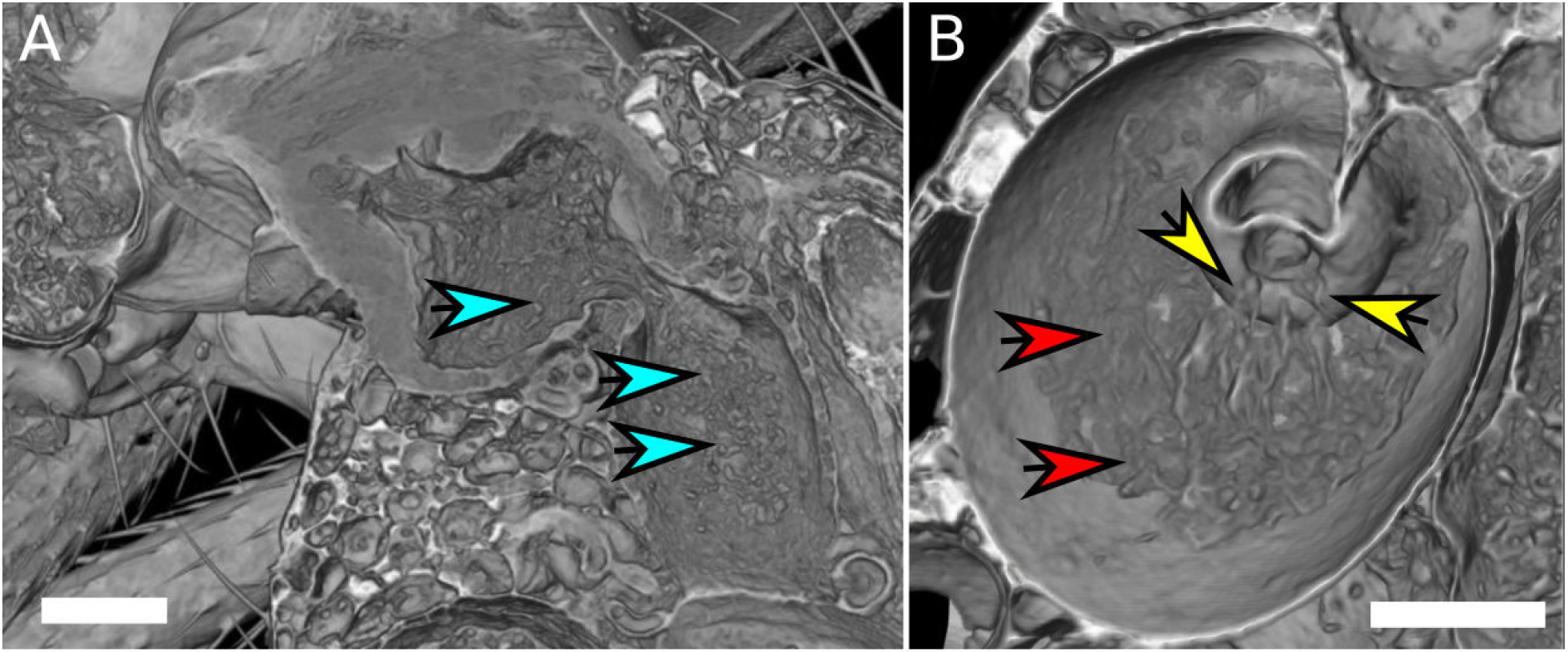
Sperm aggregates in the female reproductive tract. **A)** Female uterus harboring granular sperm aggregates, indicated by blue arrows. **B)** Spermatheca with filamentous sperm (red arrows). Filamentous sperm that enters the spermatheca is indicated by yellow arrows. The scale is 100 μm.

**Figure 5:**
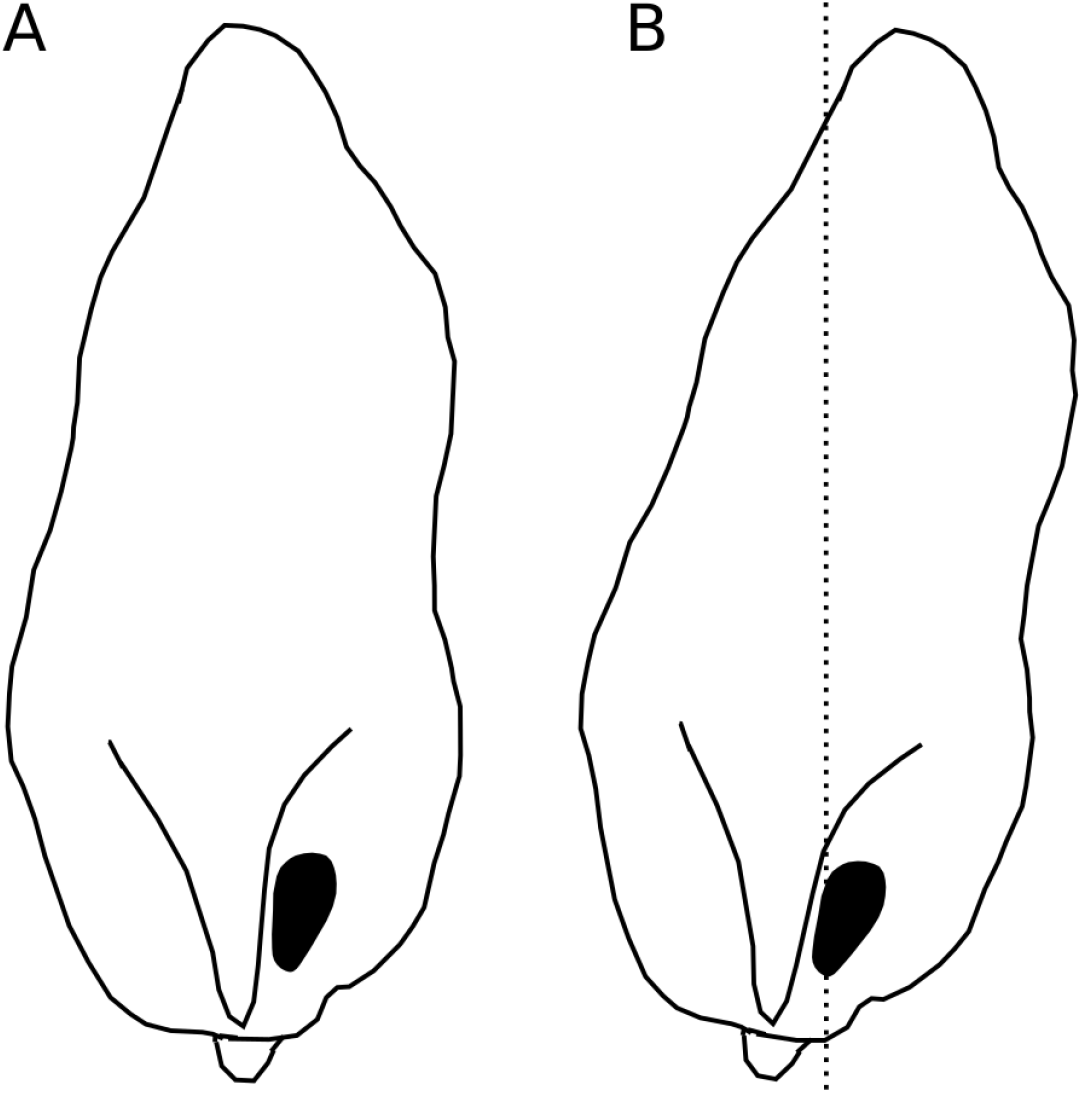
Asymmetric positioning of the male phallus. Contour drawings of the dorsal apex of the *D. pachea* male phallus. **A)** The male gonopore (filled black) is located at the right side of the phallus. **B)** The twisted complex of female and male genitalia (Rhebergen et al., 2016) is expected to shift the male apex by 6-8° which should result in a medial positioning of the male gonopore with respect to the female antero-posterior midline (dashed line).

## Discussion

### Genital asymmetry is not reflected in sperm allocation

Although we were unable to precisely quantify sperm, we observed no left or right-sided bias in the way sperm were deposited into the female reproductive tract. The distance between the phallic gonopore and the female receptacles remains large at all stages of copulation, and it appears impossible for the asymmetric phallus to preferentially deposit sperm unilaterally in the female. Sperm allocation, may therefore not be controlled by the intermittent phallus morphology, but by female structures or by components of the male ejaculate. Copulatory and post-copulatory uterus rearrangements in *D. melanogaster* females are induced by male accessory gland proteins and are associated with sperm storage and fertilization (Adams & Wolfner, 2007; Mattei et al., 2015). Selective sperm storage has been associated with female muscular activity. Storage in female yellow dung flies *Scatophaga stercoraria* was found to be sensisitve to CO_2_ anesthetization and was proposed to rely on active female muscular movements (Hellriegel & Bernasconi, 2000). In a specific case, external and non-intermittent male genital asymmetry was found to indirectly stimulate postcopulatory sperm distribution in the female in the fly *Dryomyza anilis*. During copulation, the male taps the female external genitalia after insemination with a pair of genital claspers, which can be symmetric or asymmetric. Male tapping affects sperm distribution in female sperm storage organs (Otronen & Siva-Jothy, 1991; Otronen, 1997) and males with asymmetric small claspers reveal higher fertilization success compared to males with large symmetric lobes since asymmetric male tapping probably affects sperm distribution in the spermathecae (Otronen, 1998). However, we know of no cases where the male phallus morphology influences sperm storage patterns in fly species. Phallus asymmetry in *D. pachea*, and male genital asymmetry in general, may therefore rather influence the efficiency of ejaculate transfer and stabilize the complex of male and female genitalia during copulation, but appears to be unrelated to postcopulatory sperm storage.

### Phallus asymmetry is associated with a specific orientation inside the female uterus

We identifed a curious orientation of the *D. pachea* male phallus inside the female uterus, which is strikingly different from previous findings in *D. melanogaster*. While in *D. melanogaster* and closely related species, the phallus points anteriorly towards uterus and spermathecae (Yassin & Orgogozo, 2013; Mattei et al., 2015), it points to the caudal female body wall in *D. pachea*. The *D. pachea* phallus is curved at its base. It appears that the phallus functions as a mechanical basis to anchor the male abdomen on top of the female. Sperm is released from this most caudal position of the phallus and needs to pass the phallus base to reach the anterior uterus and female sperm storage organs. This is an intriguing observation since direct ejection of ejaculate into more anterior parts of the uterus would seem more appropriate for efficient gamete transfer. The copulation complex in *D. pachea* is shifted 6-8° to the right side with respect to the female antero-posterior midline (Rhebergen et al., 2016). Extrapolating this shift to the dorsal apex of the male phallus, the right-sided gonopore is located in a medial position (Figure 5). The dorsally pointed phallus structures stretch the dorsal vagina (like a covering sheet). The ventrally pointing asymmetric phallus spurs (Acurio et al., 2019) must be directed towards the caudal body wall and can potentially function as a grasping device to ensure an internal grip of the phallus. Alternatively, they may provide ventral flaps that avoid a close covering of the ventral vagina and therefore may stabilize a ridge in which ejaculate may be transported. Overall, the asymmetric phallus morphology may function to stretch the vagina in a specific configuration to enable efficient sperm release.

### The uterus reshapes during sperm transfer

The uterus assumes a different shape during copulation compared to a virgin uterus, similar to previous observations in *D. melanogaster* (Adams & Wolfner, 2007; Mattei et al., 2015). However, broadening of the anterior part of the uterus was only observed in *D. pachea*. While it enlarges, the constriction at the junction to the spermathecal tubes remains narrow. This could potentially induce a vacuum and could either induce transport of the egg or perhaps result into a suction force to transport ejaculate towards the anterior part of the uterus. However, we have no data on the specific temporal dynamics of this re-arrangement. Future studies must analyze uterus shapes with progression of sperm and the eggs. However, *D. pachea* is rarely observed to lay eggs after a single copulation (Pitnick, 1993; Lefèvre et al., 2021), indicating that uterus reshaping might be related to other processes.

### Low efficiency transfer of a complex ejaculate mass

Sperm transport into female spermathecae was observed only in a minority of mating trials. One possible explanation for this low rate is that our laboratory conditions may not provide optimal ecological parameters for this species and this may influence copulatory processes. Also, we use a laboratory stock that has been maintained for > 20 years in captivity and which might reveal different sperm transfer frequencies with respect to wild *D. pachea*. Previous reports on *D. pachea* ejaculate transfer did not report that sperm is not always released from the male phallus during copulation (Pitnick, 1993; Pitnick & Markow, 1994). Nevertheless, our observations might indicate that ejaculation is, in fact, rare in *D. pachea*. Males of *D. pachea* were reported to transfer only 44 ± 6 sperm cells per copulation (Pitnick & Markow, 1994), which was found to be far below the female sperm storage capacity. Males produce giant sperm of >16 mm length (Pitnick & Markow, 1994) and the male investment into spermatogenesis was argued to be high compared with other *Drosophila* species. Thus, males might be selective with regard to the females to whom they would allocate their limited amount of gametes (Pitnick & Markow, 1994). Wild-caught *D. pachea* females were reported to contain sperm from at least 3-4 males and females are known to remate with various males (Pitnick & Markow, 1994). Alternatively, sperm might be discarded or absorbed during or shortly after copulation, which could lead to dumping of sperm and might relate to female controlled cryptic female mate-choice (Eberhard, 1985). In our synchrotron X-ray tomography scans, sperm was present in different parts of the reproductive tract. Our data indicates that sperm release from the phallus and sperm transport into the spermathecae might be two independent physiological processes. The overall number of gametes in each ejaculate is low and females often remate. Thus, the relative loss of gametes per copulation is also low. Overall, the reproductive strategies for both sexes may imply a high level of promiscuity to compensate for a rather inefficient sperm transfer process during copulation.

However, when copulation does result in sperm transfer, it is already released early during copulation, from at least 2 min after the start of copulation (the entire copulation lasts up to 40 minutes). Similar results were reported from remating experiments in *D. melanogaster* (Gilchrist & Partridge, 2000), where males required about 4-8 min to replace sperm from a previous mating. Intriguingly, longer copulation duration of D. *pachea*, of 18 min and 20 min yielded fewer cases of sperm transfer compared to shorter copulation durations of 8 min or 12 min. Previous analyses also revealed that copulation duration was negatively associated with sperm transfer rates in *D. pachea* (Lefèvre et al., 2021). Male dependent sperm allocation is not expected to show this pattern and was not observed in our X-ray computed tomography scans where the majority of scans revealed sperm presence in other parts of the female reproductive tract. Our observations indicate that sperm can be dumped by the female and this might occur if copulation takes too long, which can be a parameter of male performance during copulation (Eberhard, 1994). Alternatively, it could be argued that our data for the longer copulation durations are biased towards couples where the males continued copulating *because* they had not yet been able to transfer ejaculate. However, the investigated time-points were below the maximum copulation duration of 30-40 min seen in this species (Pitnick, 1993; Lang & Orgogozo, 2012; Rhebergen et al., 2016; Acurio et al., 2019; Lefèvre et al., 2021). We did not observe any couple that ended copulation before they were artificially separated.

Sperm was observed to be differentially organized in different organs. In the medial part of the uterus it had a granular shape while it was filamentous when entering the spermathecae. In *D. melanogaster*, sperm was reported to assume a torus shape inside the female uterus and such tori slowly rotate (Adams & Wolfner, 2007). It remains to be investigated at better resolution, if granular shaped sperm in *D. pachea* also approximates these tori conformations. What also need further elaboration is whether these giant sperm are motile or are transported passively by other male ejaculate components or by female peristalsis. The aggregation changes might be related to the organization of the sperm tail, which may be non- or less motile. The filament perhaps undergoes dynamic aggregations or crosslinks with other sperm tails or with different parts of the tail. This could result in compaction and elongation of sperm mass and may explain the different aggregation states observed in our microCT and synchrotron X-ray tomographic scans.

## Conclusions

The positioning of the asymmetric *D. pachea* male phallus was evaluated during copulation inside the female reproductive tract. The twisted copulation complex likely positions the right sided gonopore into a medial location at the caudal end of the vagina. This particular positioning of the male gonopore, the inflation of the oviduct during copulation and the low frequency of sperm transfer in different trials indicate a complex and inefficient sperm transport into female sperm storage organs during and after copulation. No evidence was obtained that the asymmetric male genitalia allow asymmetric sperm deposition inside the female reproductive tract. Future investigations must unravel if transfer of giant sperm might be non-autonomous in *D. pachea*, potentially mediated by re-arrangements of the female uterus and by different aggregation states of sperm during transport through the female reproductive tract.

## Acknowledgements

We would like to thank Maxi Richmond Polihronakis for discussions and for help with copulation complex preparation in *D. pachea*. We would also like to thank Kees Koops for his help and insight in starting and maintaining the fly population. We acknowledge the Paul Scherrer Institut, Villigen, Switzerland, for provision of synchrotron radiation beamtime at the TOMCAT (X02DA) beamline of the Swiss Light Source to MS and MR. We thank Dr. Federica Marone for supervision and assistance during the synchrotron experiments. Lastly, we would like to thank Rob Langelaan and Bertie-Joan van Heuven for their help during the preparation and working with the μCT-scanner at Naturalis Biodiversity Center. A part of this work was also supported by a grant of the Agence Nationale de la recherche [ANR-20-CE13-0006] given to ML.

## Supplementary Information

Data supporting this article has been deposited at the DRYAD database (https://doi.org/10.5061/dryad.5x69p8d4z).

**Table S1:** Summary table of microCT and synchrotron X-ray tomographic scans and organ size measurements.

**Table S2:** Summary table of copulation trials to test for presence of sperm mass in the female spermathecae.

**Table S3:** Sperm counts in the female reproductive tract based on synchrotron X-ray tomographic scans.

